# The Arabidopsis pattern recognition receptor EFR enhances fire blight resistance in apple

**DOI:** 10.1101/2021.01.22.427734

**Authors:** Stefano Piazza, Manuela Campa, Valerio Pompili, Lorenza Dalla Costa, Umberta Salvagnin, Vladimir Nekrasov, Cyril Zipfel, Mickael Malnoy

## Abstract

Fire blight disease, caused by the bacterium *Erwinia amylovora* (*E. amylovora*), is responsible for substantial losses in cultivated apple worldwide. An important mechanism of plant immunity is based on the recognition of conserved microbial molecules, named pathogen- or microbe-associated molecular patterns (PAMPs or MAMPs), through pattern recognition receptors (PRRs), leading to pattern-triggered immunity (PTI). The interspecies transfer of PRRs represents a promising strategy to engineer broad spectrum and durable disease resistance in crops. EFR, the *Arabidopsis thaliana* PRR for the PAMP elf18 derived from the elongation factor thermal unstable (EF-Tu) proved to be effective in improving bacterial resistance when expressed into Solanaceae and other plant species,. In this study, we tested whether EFR can affect the interaction of apple with *E. amylovora* by its ectopic expression in the susceptible apple rootstock M.26. Stable *EFR* expression led to the activation of PAMP-triggered immune response in apple leaves upon treatment with supernatant of *E. amylovora*, as measured by production of reactive oxygen species and the induction of known defense genes. The amount of tissue necrosis associated with *E. amylovora* infection was significantly reduced in the *EFR* transgenic rootstock compared to the wild-type. Our results show that the expression of *EFR* in apple rootstock may be a valuable biotechnology strategy to improve the resistance of apple to fire blight.

## Introduction

Despite being constantly exposed to a wide range of pathogens in their immediate environment, plants are resistant to most microbes. Each plant cell can trigger an immune response autonomously by employing pattern recognition receptors (PRRs) for sensitive and rapid detection of potential threats caused by pests or pathogens (Zipfel 2014). This mechanism is based on the recognition of conserved microbial molecules (pathogen- or microbial-associated molecular patterns, PAMPs or MAMPs) and is known as pattern-triggered immunity (PTI). PAMPs are important for microbes (both pathogenic and non-pathogenic), and are widely conserved across microorganisms. When these molecules are recognized by plants, a series of defense reactions are induced such as reactive oxygen species (ROS) production, callose deposition, expression of defense genes and activation of mitogen-activated protein kinase (MAPK) cascades (Wu and Zhou 2013; Lu *et al.*, 2014). Several PAMPs have been described and their recognition investigated in plants (Schwessinger and Ronald, 2012; Kawano and Shimamoto, 2013; Wu and Zhu, 2013; Boutrot & Zipfel, 2017) such as beta-glucan (Klarynski *et al.*, 2000), chitin (Kaku *et al.*, 2006), flagellin (Felix *et al.*, 1999) and the elongation factor thermal unstable (EF-Tu) (Kunze *et al.*, 2004).

EF-Tu is one of the most abundant bacterial proteins and acts in *Arabidopsis thaliana* (hereafter, *Arabidopsis*) as a very potent PAMP (Kunze *et al.*, 2004). *Arabidopsis* recognizes the *N*-terminus of the protein and specifically the *N*-acetylated peptide comprising the first 18 amino acids of EF-Tu, called elf18. EF-Tu is recognized by EF-TU RECEPTOR (EFR), a *Brassicaceae*-specific PRR (Zipfel *et al.*, 2006). EFR is related to FLAGELLIN SENSING 2 (FLS2), the receptor for the flagellin-derived PAMP flg22, and belongs to the leucine-rich repeat receptor kinase (LRR-RK) XII sub-family. Elf18 perception by EFR is specific to the *Brassicaceae* family. However, expression of the Arabidopsis *EFR* gene in *Nicotiana benthamiana* and in tomato conferred elf18 responsiveness and increased anti-bacterial resistance to these species, indicating that the downstream elements of PRR activation are conserved between Arabidopsis and other plants (Lacombe *et al.* 2010; Kunwar *et al*., 2018). Moreover, expression of *EFR* has been also reported to increase resistance to bacterial diseases in other plants such as *Medicago truncatula* (Pfeilmeier *et al.,* 2019), potato (Boschi *et al.*, 2017), rice (Schwessinger *et al.*, 2015; Lu *et al.*, 2014), and wheat (Schoonbeek *et al.*, 2015).

Fire blight, caused by the Gram-negative bacterium *Erwinia amylovora* (*E. amylovora*), is a destructive disease that strongly affects the *Rosaceae* family, including the economically important species apple and pear (Malnoy *et al.*, 2012). *E. amylovora* is classified as a member of the *Enterobacteriaceae* family, and, as such, is closely related to many important human and animal pathogens, such as *Escherichia coli, Salmonella* spp *., Shigella* spp. and *Yersinia* spp.

Control of the fire blight pathogen is difficult and is typically achieved through the eradication of entire trees displaying symptoms or via preventive spraying of copper compounds or antibiotics. However, these methods are not completely effective when used in isolation (Malnoy *et al.* 2012), and the application of antibiotics is not authorized in some countries due to the evolution of antibiotic resistance in populations of plant pathogens (Sundin *et al.* 2006). Within this framework, classical breeding and genetic engineering have been widely applied for the production of resistant cultivars and for targeted bacterial disease management (Emeriewen *et al.*, 2014a/b, 2017; Gardiner *et al.*, 2012; Malnoy *et al.* 2006, 2008, Kost *et al.*, 2015; Broggini *et al.*, 2014). However, on the one hand classical breeding is time-consuming and the quality of the obtained new cultivars is often inferior to that of commercially available apples, thus hindering consumer acceptance. On the other hand, the use of genetically modified plants is still forbidden in many countries worldwide. Due to its destructive character and to the lack of effective control mechanisms, *E. amylovora* is capable of dispersing rapidly within susceptible plants and between trees in orchards, which could result in great economic losses.

In this study, we generated transgenic apple plants expressing the *AtEFR* gene under the control of the *AtFLS2* promoter in the susceptible apple rootstock M.26. Transgenic plants were characterized by molecular and phenotypic analyses to evaluate EFR functionality and its ability to confer increased resistance to *E. amylovora*.

## Materials and Methods

### Evaluation of elf18 peptides

Nineteen elf18 peptide sequences were retrieved from Lacombe *et al*. (2010) and from the *E*. *amylovora* genome. Phylogenetic trees (maximum likelihood model) for these sequences were constructed using MEGA7 software (Kumar et al. 2006) after sequence alignment via clustalW (Larkin et al. 2007).

### Plant materials and growth conditions

For apple, wild-type and transgenic plantlets of the rootstock M.26 were grown *in vitro* following the procedure described by Malnoy *et al.* (2007), and acclimated to soil according to Bolar *et al.* (1998). *In vitro* and glasshouse plant material was cultivated with a 16/8-h light/dark period at 20-24 °C.

*Arabidopsis thaliana* plantlets of ecotype Columbia-0 (Col-0) and *efr* mutant lines were grown in soil as one plant per pot or in plates containing half MS salts medium, 1% sucrose and 1% plant agar, at 20-22,°C with a 16/8-h light/dark period.

### Plant transformation and evaluation of ploidy level

The binary vector pBin19 used for plant transformation carries the *AtEFR* open reading frame flanked by the *Arabidopsis FLS2* promoter and the *octopine synthase* terminator (pBin19/FLS2p:EFR-FLAG). This vector was transferred into *Agrobacterium tumefaciens* strain EHA105 by electroporation and used to transform the apple rootstock M.26.

Plant transformation, regeneration, propagation and acclimatization were carried out as described by Malnoy *et al.* (2006). Regenerated transgenic plants were confirmed by PCR with primers efrF (5’-CTTGAATTTATTGGGGCTGTGGCG-3’) and efrR (5’-CCTGCAAGTTCAAAAGCTTCCCGA-3’) for the presence of the transgene. Following transformation, the ploidy level of the transgenic and untransformed clones was also estimated by flow cytometry as described by Malnoy *et al.* (2005).

### Preparation of the bacterial supernatant

Overnight culture of *E. amylovora* (strains Ea273 and Ea1430) was centrifuged at 4000 *g* for 5 minutes. The obtained pellet was suspended in 10 mM phosphate buffer (pH 7,8) and boiled at 100 °C for 15 minutes. The bacterial suspension was then centrifuged at 4000 g for 5 minutes and the supernatant was collected for further experiments.

### Reactive oxygen species measurement

Leaves of *Arabidopsis thaliana* Col-0 plants or *efr* mutants were treated with the supernatant of two strains of *E. amylovora* (strains Ea273 and Ea1430). ROS measurement was performed as previously described by Zipfel *et al*. (2006).

The detection of superoxide anions in apple leaves was performed by staining the leaves with nitroblue tetrazolium (NBT). The youngest leaves of actively growing plants (control and transgenic lines) were transferred to NBT staining solution containing 5 mM NBT in 20 mM phosphate buffer, pH 6,5 (Muller *et al.*, 2009). The leaves were incubated in staining solution under constant vacuum until dark color appeared on the surfaces of the stained leaves. Stained leaves were washed with 100 % ethanol for a better visualization of the staining.

### Quantification of nptII copy number by Taqman real-time PCR

The quantification of the *nptII* copy number was used to calculate the number of T-DNA insertion events in the transformed apple plants (Supplemental Data 1) according to the Taqman real-time PCR method developed by Dalla Costa *et al.* (2019). Primers and probes used in the analysis are listed in Supplemental Table 1.

### Determination of fire blight resistance

The evaluation for fire blight symptoms was carried out in three independent experiments, as described by Malnoy *et al.* (2008). In brief, 20 vigorously growing shoots of M.26 transgenic plants and untransformed control plants (M.26 susceptible and M7 resistant cultivars) were inoculated by cutting the two youngest expanded leaves with scissors previously dipped in an *E. amylovora* suspension (strain Ea273, 5 10^7^ cfu mL^−1^). Only actively growing plants that showed a shoot length of at least 13.0 cm were considered for the experiments. Disease severity was rated 1 month after inoculation as percentage of the length of the necrosis/total length of the shoot growth, as described by Campa *et al.,* 2018. Data interpretation and statistical analysis were performed according to Pompili *et al*., 2020.

Inoculated shoot tips became orange-brown and eventually dark brown, often with production of cloudy ooze droplets which darkened with time. These symptoms proceeded basipetally on the inoculated shoot, and eventually terminated depending on the susceptibility of the shoot resulting from its genetics and the vigor of its growth. Three to fifteen biological replicates for each plant line were inoculated with *E. amylovora* strain Ea273 (10^7^ cfu mL^−1^) in each of the three independent experiments performed. Only actively growing plants that showed a shoot length of at least 13 cm were considered for the experiments. Data collection was performed according to Campa *et al*. (2018). Statistical analysis was performed using the DellTM StatisticaTM Software version 13.1. As the three experiments showed the same trend, measures of each plant line were merged and analysed as a single experiment. Nonparametric Kruskal–Wallis test was used to compare groups as data did not show a normal distribution. Subsequently, all groups were compared simultaneously by multiple comparisons of mean rank. Statistical analysis was performed with probability level (alfa) = 0.05.

#### Bacterial concentration in plants

Fifteen days post inoculation, three sections of inoculated leaf or healthy tissue from *EFR* transgenic plants and control M.26 were disrupted with two metal balls (5 mm) in 500 mL King B media in a GenoGrinder (2 9 20 s, 1250 spm) to release bacteria from the leaf apoplast. A 10^−1^–10^−6^ dilution series was made in LB medium and 100 μL from each dilution were spotted on Kado media and incubated at 28 °C for 24 h. The number of colonies was counted to estimate the number of colony forming units (cfus) per inoculation site. Four independent experiments were performed.

#### Total RNA isolation and gene expression analysis

Total RNA was extracted from apple leaves with the Spectrum Plant Total RNA Kit (Sigma-Aldrich, St. Louis, MO). Two micrograms of total RNA were treated with DNAse (Ambion) and then reverse transcribed with SuperScript VILO cDNA SynthesisKit (Invitrogen, Carlsbad, CA), according to manufacturer instructions. Real time quantitative PCR was conducted using Biorad CFX96 System (Bio-Rad Laboratories, Hercules, CA) as described in Campa *et al.* (2019). Each reaction was performed in triplicate. Gene-specific primers are reported in Table S1. Apple *Actin* (*actin2*), *Ubiquitin* (*Ubi*) and *Elongation Factor 1* (*EF1*) were used as internal reference genes for normalization (Table S1). The Biorad CFX manager software version 3.0 was used to analyse the relative expression level. Statistical analysis was performed using Statistica software version 9.1 (StatSoft Inc., 2010). Gene expression variance was analysed by One-way ANOVA followed by post hoc Tukey’s test. All analyses were performed using three biological replicates.

## Results

### Evaluation of E. amylovora EF-Tu eliciting activity

Based on the EF-Tu sequence from *Escherichia coli* (*E. coli*) (Kunze *et al.* 2004), the sequence of elf18 was retrieved from different *Erwinia* species (Figure 1A,B). Alignment of the *Erwina* sequences with 14 selected elf18 sequences revealed that all of the elf18 sequences derived from *Erwinia* species are grouping with the *E. coli* elf18 sequence (Figure 1 B). A strong conservation of the consensus in the elf18 peptides can be observed (Figure 1A). Indeed, 9 of the 18 amino acids of elf18 conserved domains are preserved in all the bacteria species (Figure 1A), while specific amino acid substitution allows to define specific clades of bacteria (Figure 1B).

**Figure 1:**
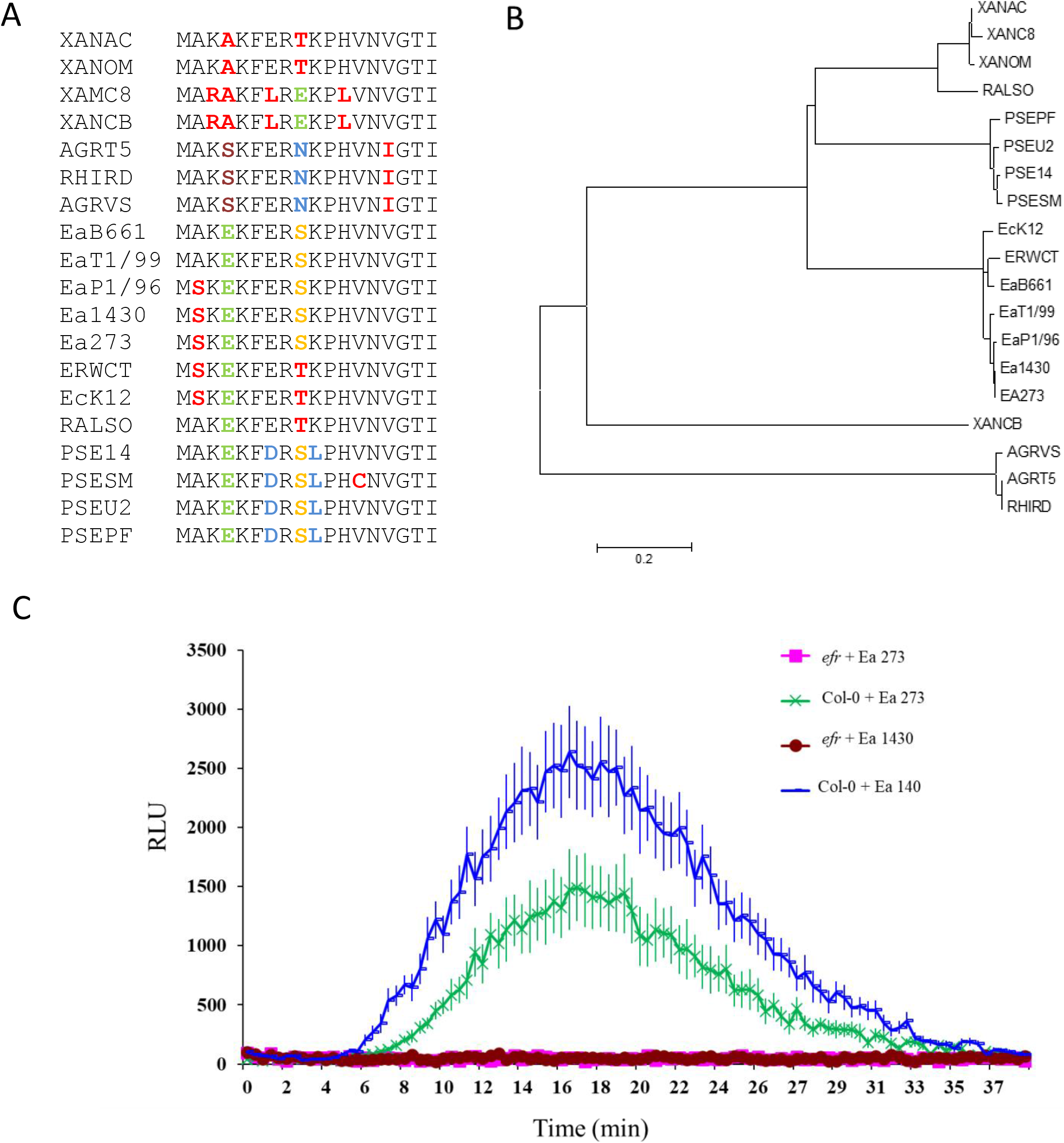
Conservation of elf18 epitope in *E. amylovora* A) Alignment of elf18 regions from selected bacteria: XANAC: *Xanthomonas Axonopodis pv. citrus,* XANOM: *Xanthomonas oryzea pv. oryzea;* XANC8: *Xanthomonas Campestris pv. Campestris 8004*; XANCB: *Xanthomonas Campestris pv. Campestris B100;* AGRT5: *Agrobacterium tumefaciens C58*; RHIRD: *Agrobacterium tumefaciens; AGRVS: Agrobacterium vitis S4*; EaB661: *Erwinia billingeae 661*; EaT1/99: *Erwinia tosmaniensis 1/99; Ea1430: Erwinia amylovora 1430*; Ea273: *Erwinia amylovora 273;* EP1/79*: Erwinia pyrifoliae Ep1/79*; ERWCT: *Erwinia carotovora ssp. Atroseptical/P. Atrosepticum;* EcK12: *Echerichia coli K12;* RALSO: *Ralsonia solanacearum* GMI1000; PSE14: *Pseudomonas syringae pv. Phaseolicola* 1448; PSESM: *Pseudomonas syringae pv. tomato* DC3000; PSEU2: *Pseudomonas syringae pv. syringae* B728a; PSEPF: *Pseudomonas fluorescence* pfo-1. B) Phylogenetic tree of the elf18 sequences from the selected bacteria mentioned above. C) Reactive oxygen species generation in *Arabidopsis thaliana* wild-type and *efr* mutant in response to mock or *Erwinia amylovora* (Ea1430 or Ea273) supernatant treatments measured in relative light units (RLU). Data are average ± SE (n=10). This experiment was repeated 2 times with similar results.

To verify the activity of *E. amylovora* EF-Tu elicitor, heat-killed *E. amylovora* extracts were applied to wild-type and *efr* mutant *A. thaliana* leaf discs. Extracts from *E. amylovora* strains were able to elicit ROS production in wild-type Arabidopsis leaves; in contrast, this response was abolished in *efr* mutant leaves (Figure 1 C). These results indicate that the main elicitor present in the extract derives from EF-Tu.

### Generation of transgenic apple, molecular evaluation of the transgenic events and ploidy level

The susceptible apple rootstock M.26 was transformed to express *EFR* (see methods for details). A total of 6 transgenic lines were obtained from one transformation experiment. All the transgenic lines were confirmed by PCR for the presence of the *EFR* gene and then characterized for the T-DNA integration copy number into the genome. The lines TM-2017, TM2108 and TM2114 had one T-DNA insertion, while the lines TM2111, TM2113 and TM2112 had 3, 5 and 8 copies of inserted T-DNA, respectively. Moreover, the ploidy level determined by flow cytometry showed that all of the transformed lines were diploid, like the non-transformed M.26. Thus, expression of *EFR* was evaluated in plants grown at greenhouse conditions. RT-qPCR analysis showed that *EFR* was expressed in leaves of all the transgenic lines before *E. amylovora* infection and significantly increased 10- to 20-fold 24 h post-infiltration with *E. amylovora* bacterial suspension or supernatant, respectively (Figure 2).

**Figure 2:**
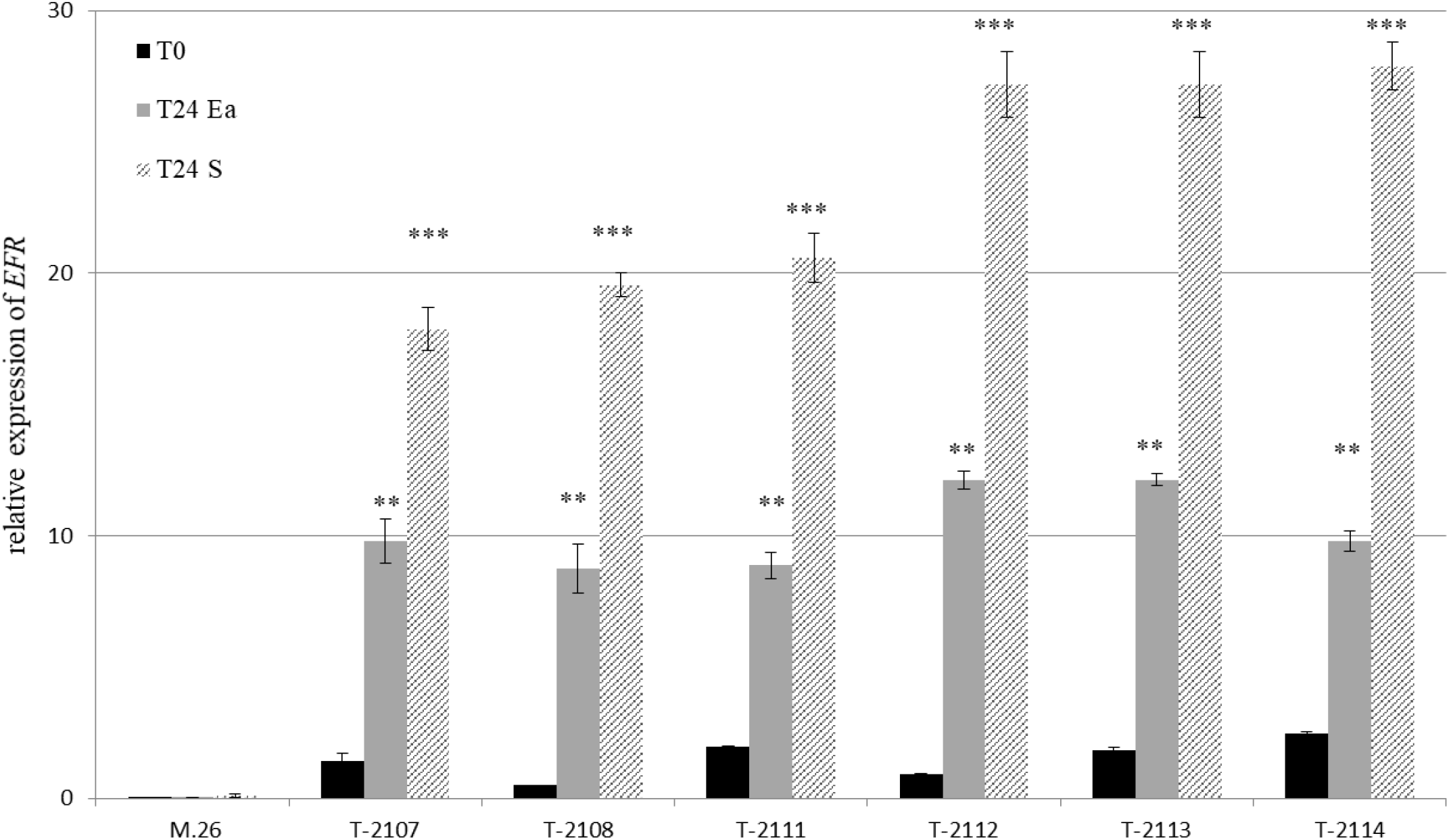
Expression pattern of *EFR* in transgenic apple rootstock M.26 compared to wild-type before and 24 h after treatment with supernatant of *E. amylovora* (S) or with *E. amylovora* (Ea) at 10^7^ cfu mL^−1^. This experiment was repeated 3 times with similar results. Bars indicate the mean values ± SE. Considering mock- and pathogen-treatments separately, asterisks indicate statistically significant differences of datasets from the corresponding dataset at time zero (0 h), according to one-way ANOVA followed by post hoc Dunnett’s test (α = 0.05).

### EFR expression in apple induces oxidative burst and defense gene upon Erwinia inoculation

To test the effect of the expression of *EFR* on the production of ROS in the transgenic lines (untreated and treated with *E. amylovora* or supernatant of *E. amylovora*), the production of superoxide anions was examined using NBT staining in those lines showing only one insertion copy. As shown in Figure 3A, 48 h after infiltration with bacteria or supernatant of *E. amylovora* (strain Ea273), a significant accumulation of superoxide anions can be observed in the two M.26 transgenic lines (TM-2107 and TM-2108) tested. In the control M26, a small superoxide anion staining can be noticed after treatment with *E. amylovara* supernatant compared with infection with *E amylovora*. In the M.26 *EFR* transgenic lines a low superoxide anion staining can be observed after water treatment, demonstrating that ROS burst is effectively due to the induction of *EFR* after treatments.

**Figure 3:**
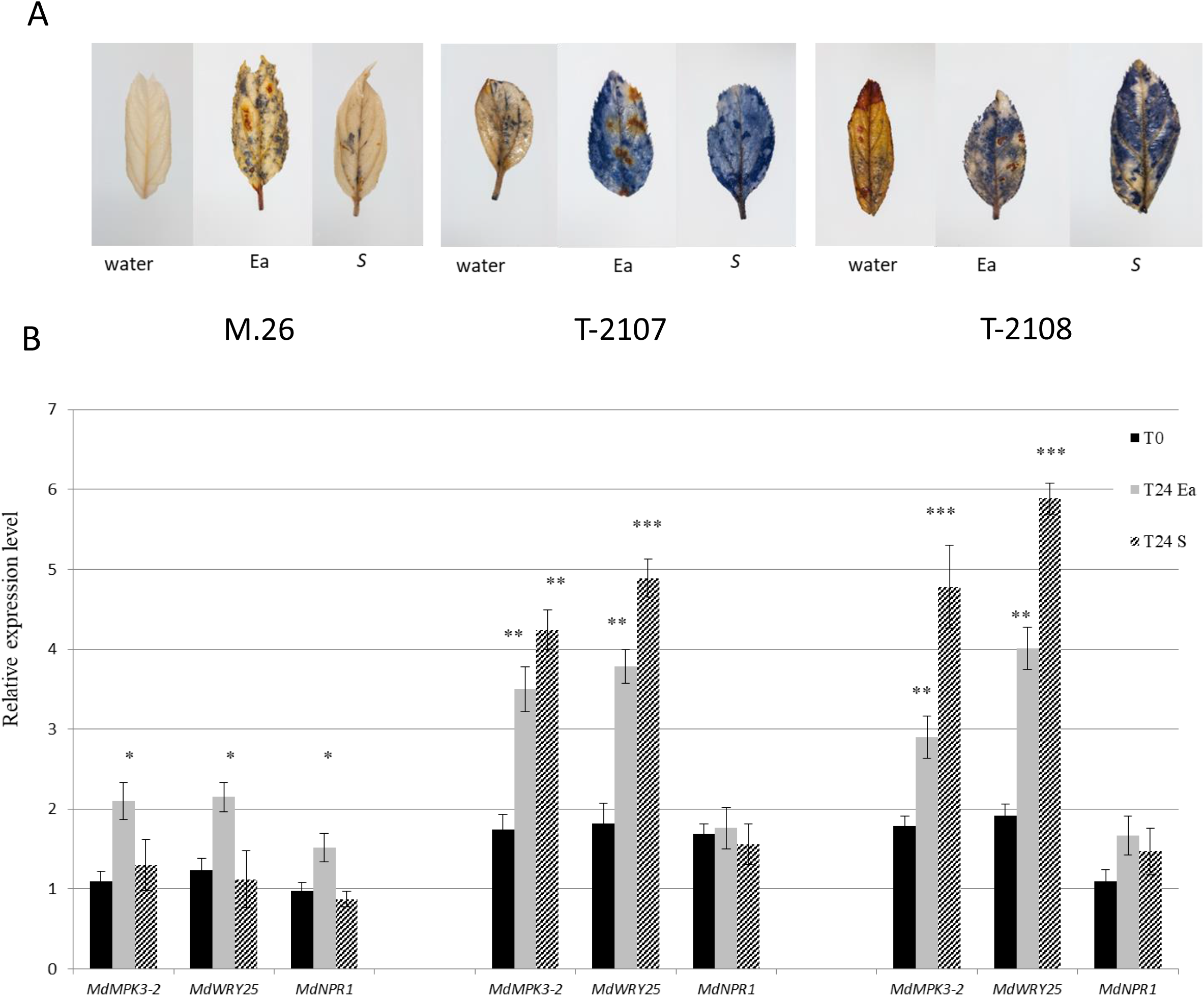
Evaluation of ROS production (A) and transcript levels of defense genes (B) in *EFR* apple transgenic lines before and 24 h after treatment with supernatant of *E. amylovora* (S) or with *E. amylovora* (Ea) at 10^7^ cfu mL^−1^. This experiment was repeated 3 times with similar results. Bars indicate the mean values ± SE. Considering mock- and pathogen-treatments separately, asterisks indicate statistically significant differences of datasets from the corresponding dataset at time zero (T0), according to one-way ANOVA followed by post hoc Dunnett’s test (α = 0.05).

PAMPs elicit the rapid and strong expression of *MPK3*, *MPK6* and other defense genes in *Arabidopsis* (Pitzschke *et al.* 2009). This observation led us to examine whether ROS marker genes were induced in our transgenic lines after bacterial infection. Transcript levels of *MdMPK3-2* and *MdWRKY25* increased in the transgenic lines after infiltration with bacteria or supernatant of *E. amylovora* (strain Ea273), whereas a small increase can be observed in M.26 only after inoculation with the bacteria suspension of *E. amylovora* (Figure 3B). In contrast, no significant increase of expression can be observed for the *MdNPR1* gene in all the tested samples.

### Evaluation of transgenic lines on their resistance to fire blight

To characterize the response of the transformed lines to fire blight, 20 acclimated biological replicates per line were inoculated with *E. amylovora* Ea273 at 5 × 10^7^ cfu mL^−1^ in three independent experiments. The response was similar in the replicated experiments, as indicated by low confidence intervals of means. The control M.26 showed an average necrosis of 70 %, indicating its high susceptibility to *E. amylovora*, while 2 *EFR* transgenic lines showed significantly lower necrosis (Figure 4A, B) compared to the wild-type plants. Indeed, all *EFR* transgenic lines exhibited a reduction in the necrosis of 70% to 80% compared to the control, one month post-inoculation with *E. amylovora* strain Ea273. Notably, this level of resistance was comparable to what observed in the resistant genotype M.7 (Malnoy *et al.* 2007). In the transgenic lines, the necrosis due to *E. amylovora* was restricted in an area of few centimetres from the inoculation point, allowing the growth of new healthy shoot under the necrotic tissue (Figure 4A). For the control, almost all the shoots were necrotic 15 days after inoculation, without new growing shoots. Moreover, the reduction of necrosis in the transgenic lines (T-2107, T-2108 and T2112) 5 days after infection correlated with a 100-fold reduction in the amount of *E. amylovora* that could be recovered from the transgenic plants compared to control plants. Indeed, 120 h after inoculation the bacterial population, evaluated by a population dynamic study, had reached 5 × 10^8^ cfu mL^−1^ (+/− 0,5 cfu mL^−1^) in the inoculated wild type leaves and 10^5^ cfu mL^−1^ (+/− 0,7 cfu mL^−1^) in the inoculated transgenic leaves. Furthermore, no *E. amylovora* bacteria was detected in the healthy transgenic tissues compared to the control.

**Figure 4:**
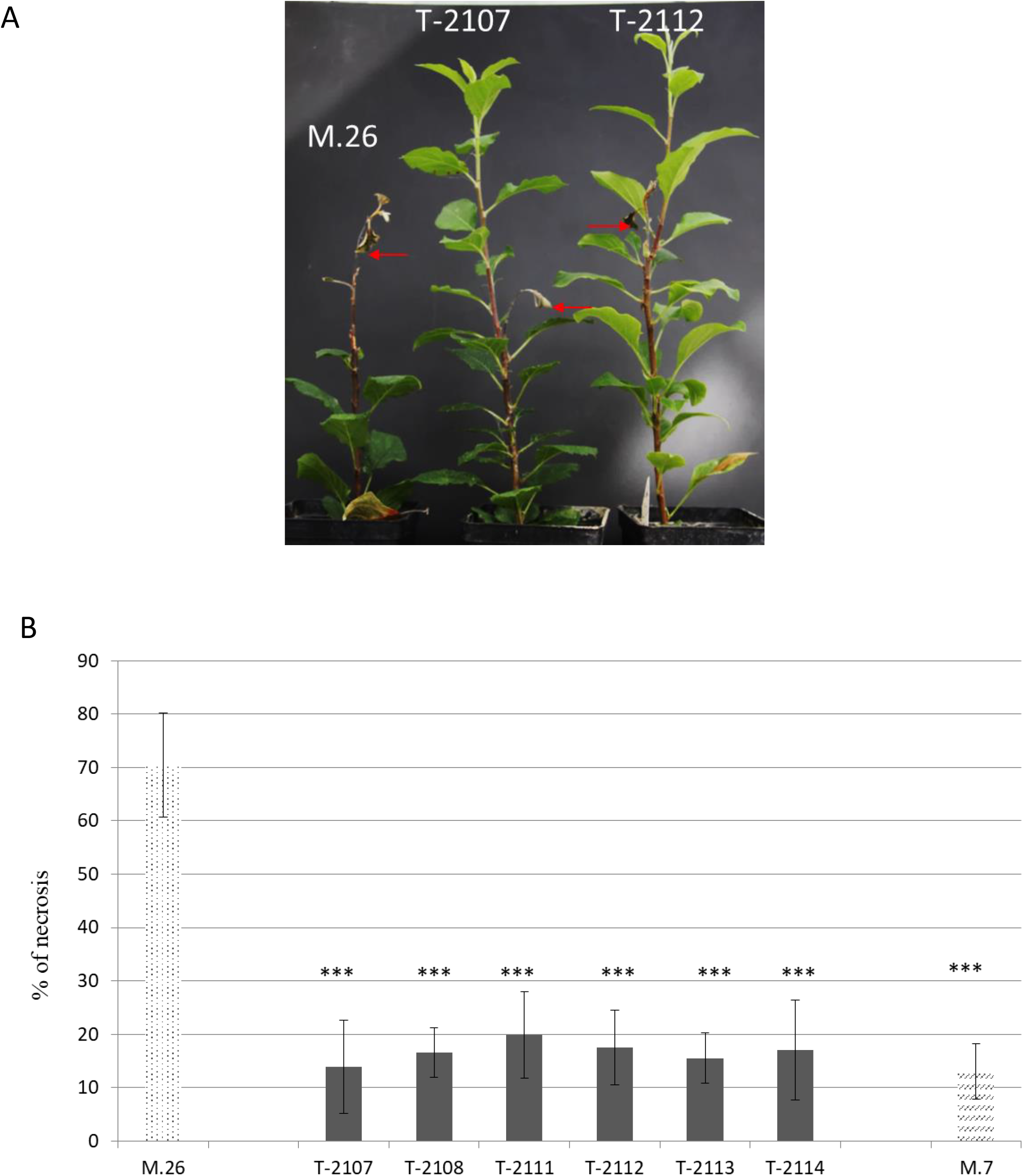
Fire blight severity in *EFR* transgenic apple rootstock M.26. A) Percentage of necrosis (shown with 95 % confidence interval bar). Susceptibility of the EFR transgenic rootstock lines to *E. amylovora* in comparison with the wild-type M.7 apple rootstock (resistant to fire blight) and M.26 (susceptible to fire blight) apple rootstock. Lengths of shoot and the portions that appeared necrotic were measured 1 month after inoculation with *E. amylovora.* Transgenic rootstock susceptibility is significantly different from M.26 at P < 0,001 (three asterisk), according to Kruskall and Wallis test. B) Pictures, taken 1 month after inoculation, showing the fire blight induced necrotic phenotype in the wild-type M.26 and in selected transgenic lines. Data are average ± SD (n=20). This experiment was repeated 3times with similar results.

## Discussion

Apple is one of the major fruit crops in the world with over 80 million tons produced each year (FAO, 2017). To maintain a high and good level of production, a considerable amount of chemical treatments is necessary (15-30 treatments/year) to protect apple against its wide range of pathogens. Major management strategies for bacterial diseases in *Malus* include biological control, cultivation practices, and breeding for disease-resistant cultivars. Although chemical control has proved to be very successful for pests and fungal diseases in apple, in the case of fire blight, no truly effective bactericide is commercially available (Malnoy *et al.*, 2012). Therefore, breeding for disease resistance has been considered as the most effective and environmentally friendly approach to control bacterial diseases in apple. However, valuable resistant germplasms within domesticated *Malus* are limited for bacterial diseases such as fire blight (Volk *et al.,* 2005 2008). Transfer of resistance genes is an alternative way to improve plant immunity and that has been already done in apple and pear to increase resistance to fire blight (Malnoy *et al.*, 2005, 2006, 2008; Broggini *et al.,* 2014). In the case of the resistance conveyed by the *Rin4* gene, it has been overcome by the pathogen over a period of time (Vogt *et al.,* 2013). In order to develop novel forms of disease control, understanding the key determinants of the relevant bacterium–host interaction is crucial (Sundin *et al.* 2016). One of these key determinants is the EF-Tu protein. Sequence alignment revealed that the key residues in the elf18 epitope of *E. amylovora* are conserved, thus suggesting that EF-Tu-derived from *E. amylovora* is recognized by *EFR* (Figure 1). In addition, we showed that supernatant of two strains of *E. amylovora* can induce ROS production in *A. thaliana* but not in the *efr* mutant.

In this study, we generated transgenic rootstock of apple that express AtEFR to confer recognition of EF-Tu from *E. amylovora*. The transgenic rootstock M.26 plants recognize the EF-Tu-derived peptide elf18, leading to the production of ROS burst in leaves as previously reported in other plants expressing EFR (Boschi *et al.* 2017). These results indicate that components required for EFR-triggered downstream immune signaling are conserved not only in *Solanaceae* but also in apple, the 50-million year divergence between *Maloideae* species and *Arabidopsis thaliana*.

We demonstrated that all the generated apple transgenic lines present a high level of resistance to fire blight similar to the resistant control (M.7) (Malnoy *et al.* 2007). Furthermore, transgenic rootstock plants expressing *EFR* did show constitutive activation of defense responses, as measured by ROS production, and expression of defense genes. They also did not exhibit any developmental or growth defects over several generations (data not shown). After treatment with supernatant of *E. amylovora* or inoculation with *E. amylovora,* a strong ROS production and increase of defense gene expression could however be observed, documenting the inducibility of EFR activation in the transgenic lines.

The use of this transgenic rootstock can be interesting for protecting apple cultivars against fire blight or other bacterial pathogens such as *Agrobacterium tumefaciens*. Indeed, rootstocks have been used in agriculture for over 2000 years in Asia to improve production, reduce disease susceptibility, and increase sustainable agriculture by reducing inputs (Haroldsend *et al.*, 2012; Kubato *et al.,* 2008). The first contribution of rootstock for food cultivation was by domestication and propagation of woody species difficult to root from cutting such as apple, pear and plum (Mudge *et al.,* 2009). Nevertheless, the best example of rootstock contribution for food security was the rescue of the European grape and wine industry from the devastating effects of the soil born insect phyloxera in the 19^th^ century (Pouget 1990).

Our transgenic rootstock can be used as resistant rootstock against fire blight in countries that already allow the cultivation of GM organisms, and might be a further motivation for the deregulation of GM crops to improve disease resistance in countries that do not do so yet. It may also improve the resistance of the scion to the disease; indeed, it was proven that grafting can be useful for the transport of signaling molecule from the rootstock to the scion to gain virus resistance (Kasai *et al.*, 2011; Zhao and Song 2014).

## Acknowledgements

This work was funded by the Autonomous Province of Trento. Work in the Zipfel laboratory on the interfamily transfer of EFR is funded by the Gatsby Charitable Foundation and the 2Blades Foundation.

## Author contributions

S.P. conducted the majority of experiments and wrote the manuscript. M.C., V.P., U.S and D.C.L. contributed performing the experiments, and revised the manuscript. C.Z. and M.M designed the project, contributed to designing the experiments and revised the manuscript. All authors read and approved the manuscript.

---

**Table S1:**
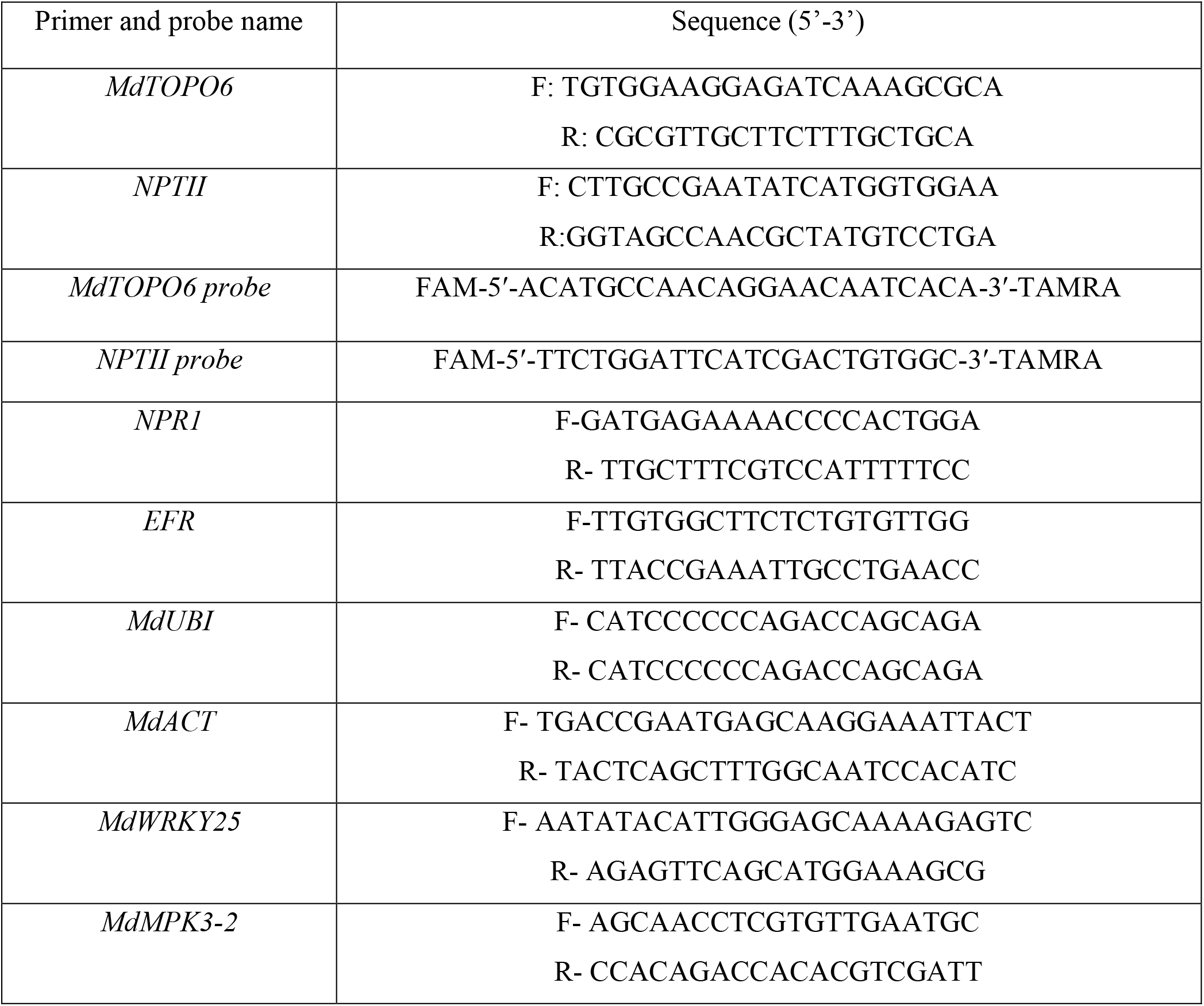
Sequences of primers and probes used for the quantification of *nptII* copy number, and primer used for QPCR.

